# Category-specific item encoding in the medial temporal lobe and beyond: The role of reward

**DOI:** 10.1101/2021.01.22.427769

**Authors:** Heidrun Schultz, Jungsun Yoo, Dar Meshi, Hauke R. Heekeren

## Abstract

Forming new memories is a fundamental part of human life, and the medial temporal lobe (MTL) is central to memory formation. Recent research suggests that within MTL, the perirhinal and parahippocampal cortices (PRC, PHC) process object and scene memory, respectively, whereas the hippocampus (HC) is agnostic to stimulus category. It is unclear, however, whether MTL category specificity extends to item encoding. Furthermore, MTL does not act in isolation: Reward-related memories are formed in interplay with the dopaminergic midbrain (substantia nigra/ventral tegmental area, SNVTA) and amygdala (AMY), but it is unclear whether reward modulates neural item encoding in a category-specific way. To address these questions, we had 39 healthy volunteers (27 for all memory-based analyses) undergo functional magnetic resonance imaging while they solved an incidental encoding task, which paired objects or scenes with high or low reward, followed by a next-day surprise recognition test. Behaviourally, high reward preferably enhanced object memory. Importantly, neural activity in PRC and PHC reflected item encoding of objects and scenes, respectively. Moreover, AMY encoding effects were selective for high-reward objects, with a similar pattern in PRC. SNVTA and HC showed no clear evidence of item encoding. The behavioural and neural asymmetry of reward-related encoding effects may be conveyed through an anterior-temporal memory system, including AMY and PRC, potentially in interplay with the ventromedial prefrontal cortex (vmPFC).

## 1 Introduction

The ability to turn experiences into new episodic memories is a central part of life. Beginning with the famous patient H.M. in the 1950s (Scoville and Milner, 1957), a large body of research indicates a critical role for the medial temporal lobe (MTL) in the encoding and retrieval of episodic memories (Zola-Morgan and Squire, 1990; Eichenbaum et al., 2007; Squire and Wixted, 2011). The MTL is not homogeneous, however, but consists of several subregions including the hippocampus (HC), perirhinal cortex (PRC), and parahippocampal cortex (PHC), which differ in their cytoarchitecture as well as anatomical connectivity with the rest of the brain (Burwell, 2000, 2001; Eichenbaum et al., 2007; van Strien et al., 2009; Ding et al., 2016; Berron et al., 2017). What, then, are the individual contributions of these subregions to episodic memory? A powerful predictor of MTL subregion function is anatomical connectivity (Davachi, 2006; Eichenbaum et al., 2007). Data from non-human primates and rodents indicate differential connectivity of the MTL input/output regions, PRC and PHC, to the ventral and dorsal visual stream, respectively (Suzuki and Amaral, 1994a; Burwell and Amaral, 1998a). Therefore, these regions are thought to process information in a category-specific way, with object-related processing in the PRC, and spatial processing in the PHC (Eichenbaum et al., 2007; Robin et al., 2018). This information is then relayed, both directly and via the entorhinal cortex (EC), to the HC, where these streams converge (Witter and Amaral, 1991; Suzuki and Amaral, 1994b; Tamamaki and Nojyo, 1995; Burwell and Amaral, 1998b; Lavenex and Amaral, 2000; Doan et al., 2019). HC’s role in memory is therefore thought to be associative and agnostic to stimulus categories (Davachi, 2006; Eichenbaum et al., 2007). In humans, functional connectivity of MTL subregions resembles these anatomical findings in animals (Kahn et al., 2008; Libby et al., 2012; Maass et al., 2015; Navarro Schröder et al., 2015). Indeed, human patient studies support the notion of a category-specific organisation of MTL subregions (Lee et al., 2005a, 2005b; Taylor et al., 2007; Mundy et al., 2013). Functional imaging studies have localised object-related and spatial processing to PRC and PHC, respectively, during a range of tasks including perception (Litman et al., 2009; Liang et al., 2013; Berron et al., 2018), associative encoding (Awipi and Davachi, 2008; Staresina et al., 2011), associative retrieval (Staresina et al., 2012, 2013; Mack and Preston, 2016; Schultz et al., 2019), working memory (Libby et al., 2014), short-term memory reactivation (Schultz et al., 2012), and recognition memory (Martin et al., 2013; Kafkas et al., 2017). It is unclear, however, whether this object-related vs. spatial distinction in PRC vs. PHC generalises to item encoding. There are numerous reports of PRC involvement in item encoding (Davachi et al., 2003; Ranganath et al., 2004; Staresina and Davachi, 2008; Wang et al., 2013). However these studies did not contrast categories; hence it is unclear whether these PRC item encoding effects are category-specific. On the other hand, PHC and the larger parahippocampal place area (PPA) have been implicated in category-specific item encoding for scenes compared to faces in studies that either did not report effects in PRC (Prince et al., 2009), or showed category-independent item-encoding effects in PRC for both faces and scenes (Preston et al., 2010). Given the reports outlined above that PRC and PHC respond to the viewing of object-related and spatial stimuli, and are differentially involved in their *associative* encoding, there are strong reasons to expect a similar dissociation of MTL cortices *for item encoding* of objects and scenes.

A mostly separate line of research has investigated how memories are formed in the first place, regardless of category. The dopaminergic reward system, to which the MTL is densely connected (Haber and Knutson, 2010; Shohamy and Adcock, 2010; Miendlarzewska et al., 2016), plays a key role. In a seminal article, Lisman and Grace (2005) have described a mechanism in which the HC and dopaminergic system interact to encode new long-term memories. Here, hippocampal novelty signals are relayed via the ventral striatum (VS) to the dopaminergic midbrain, where they trigger a dopamine response that in turn promotes long-term potentiation in HC (Lisman and Grace, 2005). In humans, reward and reward motivation enhance memory formation, accompanied by functional modulations of the HC and dopaminergic midbrain (substantia nigra/ventral tegmental area, SNVTA) (Wittmann et al., 2005, 2008; Adcock et al., 2006; Wolosin et al., 2012; Miendlarzewska et al., 2016). Importantly, not only the HC, but also the MTL cortex and adjacent amygdala (AMY) are innervated by the SNVTA (Beckstead et al., 1979; Scatton et al., 1980; Insausti et al., 1987; Oades and Halliday, 1987) and connected to other regions of the reward network including the ventromedial prefrontal cortex (vmPFC) (Russchen and Price, 1984; Amaral and Insausti, 1992; Carmichael and Price, 1995; McIntyre et al., 1996; Kondo et al., 2005; Price, 2007; Kondo and Witter, 2014).

However, these two lines of research – category specificity and reward enhancement of memory – have never been jointly investigated. It is therefore unclear whether reward enhances memory formation for objects and scenes in similar ways. Given the connectivity outlined above, memory formation for objects and scenes could be enhanced in a category-independent way through hippocampal mechanisms, and/or in a category-specific way through modulation of MTL cortex. The PRC and AMY may play a unique role in reward-enhanced item encoding. PRC may link object features to reward information (Miyashita, 2019), and PRC and AMY are both parts of a hypothesised “anterior temporal system” (AT) that is thought to represent the (motivational) salience of unitised entities such as objects (Ranganath and Ritchey, 2012; Ritchey et al., 2015). Indeed, it has been shown that another strong behavioural motivator – emotion – selectively enhances item encoding in PRC and AMY, but not context encoding in PHC and HC (Ritchey et al., 2019). Findings of item vs. context dissociations in MTL, in turn, may be tied to object-related vs. spatial processing (Davachi, 2006). It follows that reward modulation of neural item encoding effects may be at least in part category-specific.

Hence, we have identified two open questions. One, does category specificity in the MTL cortex extend to item encoding? Moreover, two, does reward modulate item encoding in a category-specific manner? To close these gaps in the literature, we investigated the neural effects of successful item encoding for two categories (objects and scenes), fully crossed with two reward magnitudes (high and low). Thirty-nine participants (27 for all memory-based analyses) underwent functional magnetic resonance imaging (fMRI) while they solved an incidental encoding task in which novel objects and scenes predicted high or low reward. One day later, they returned to the lab for a surprise recognition memory test. Behaviourally, we expected high reward to improve recognition memory for both objects and scenes. We furthermore expected activity in MTL, AMY, and SNVTA to reflect this enhanced encoding in a category-independent (HC, SNVTA) and category-specific manner (PRC/AMY for objects, PHC for scenes).

## 2 Materials and Methods

### 2.1 Participants

A total of 39 participants (“full sample”, 25 female, mean age 24.2 years, range 18-32) took part in the fMRI study. All were right-handed, had normal or corrected-to-normal vision, and were native speakers of German. A subsample of 27 participants (“memory sample”, 19 female, mean age 24.6 years, range 19-32) was selected for memory-based analyses (model 1) based on their memory performance (corrected recognition [CR] > 0.083 in each of the four conditions, see below; this threshold was chosen as a compromise between memory performance in the subsample and experimental power). Additional non-memory based analyses (model 2) were carried out in the full sample. All participants gave written informed consent in a manner approved by the local ethics committee. They received monetary reimbursement for their participation (€8/hour plus up to €5 reward during the incidental encoding task). Thirty-three fMRI datasets were complete, contributing 240 trials each, 6 suffered partial data loss due to equipment malfunction, contributing 200-238 trials each.

### 2.2 Stimuli and Procedure

A total of 360 colour photographs of objects and scenes (180 each) were obtained from established databases (Brady et al., 2008; Konkle et al., 2010a, 2010b) and an internet search. Of these, 240 (120 objects, 120 scenes) were used as targets in the incidental encoding task (Figure 1A), the others served as distractors in the surprise recognition task (Figure 1B). Assignment of images to targets and distractors was randomised for each participant. An additional 8 photographs (4 objects, 4 scenes), not included in the 360 experimental stimuli, were obtained from the same sources and used during training before the incidental encoding task (see below). Each image was sized 256×256 pixels. All tasks were programmed using Presentation® software (Version 18.2, Neurobehavioral Systems, Inc., Berkeley, CA, www.neurobs.com). The fMRI task was projected onto a mirror mounted on the head coil, and responses were collected using an MRI-compatible button box. The behavioural recognition task was presented on a laptop.

**Figure 1.**
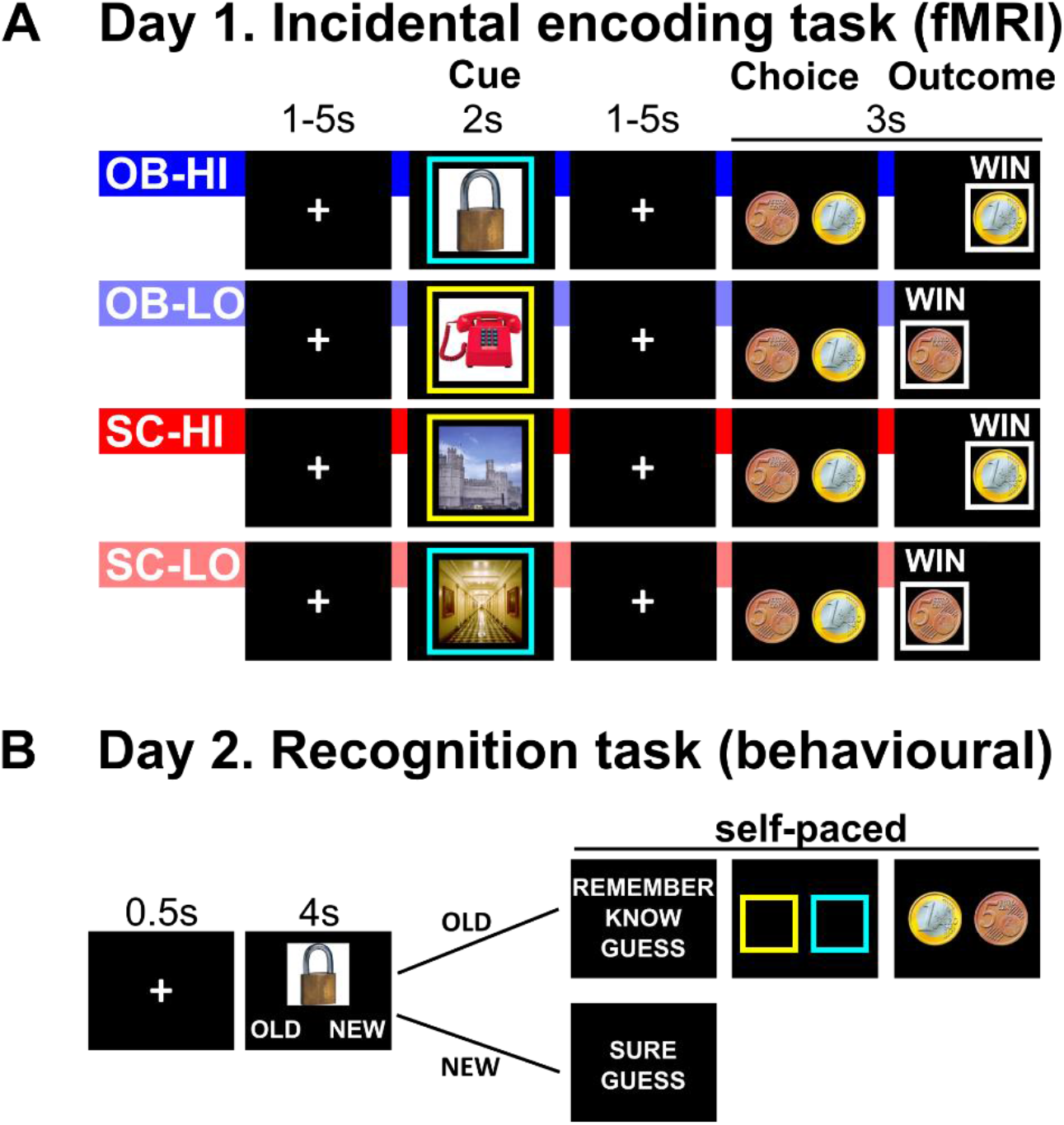
Experimental paradigm. A. Day 1 Incidental encoding task. B. Day 2 Recognition task. Note that text options (e.g. Remember – Know – Guess) were arranged horizontally in the experiment. See main text for details. Abbreviations: OB: objects, SC: scenes, HI: high reward, LO: low reward.

#### Day 1: Incidental Encoding Task (fMRI)

The incidental encoding task was designed to fully cross stimulus category (objects, scenes) and anticipated reward magnitude (high, low). This resulted in the following experimental conditions: object-high (OB-HI), object-low (OB-LO), scene-high (SC-HI), scene-low (SC-LO). The task was presented in 6 runs of 40 trials each (240 trials total), with a short break between runs. Trials were pseudo-randomised so that each run contained equal trial numbers of each condition. Additionally, no more than 3 trials belonging to the same level of each factor (category, reward) appeared in a row. Each trial consisted of a cue, choice, and outcome phase (see Figure 1A for example trials for each condition). During the *cue phase* (2s), an image (object or scene) was presented surrounded by a yellow or blue frame. Importantly, the combination of image category and frame colour coded reward magnitude: In a given run, an object with a yellow frame or a scene with a blue frame indicated high reward, while an object with a blue frame or a scene with a yellow frame indicated low reward. Category-frame combinations alternated over runs, with the run order counterbalanced over participants. The combinations were explicitly instructed at the beginning of each run (note that this is not a reward learning task). The cue phase was followed by a variable fixation (1-5s), whose duration was drawn randomly from a uniform distribution. During the following *choice phase,* two coins were presented, respectively, on the left and right side of the screen and participants were asked to indicate whether they anticipated a high (1€) or low (5C) reward in this trial. Response sides were assigned randomly. Upon button press, the chosen coin was outlined by a white frame for 0.5 s or until 2s after choice onset, whichever was shorter. In the *outcome phase,* only the chosen coin (1€ or 5C) remained on the screen, together with the word GEWINN (“Win”) if the participant had indicated the correct reward magnitude during the choice phase. For incorrect choices, the coin would be crossed out, and the word NICHTS (“nothing”) appeared on the screen. If the participant failed to press the button, the coin would also be crossed out, and the words ZU LANGSAM (“too slow”) appeared on the screen. The outcome phase lasted 1s or until 3s after choice onset, whichever was longer. Trials were offset by a variable fixation interval of 1-5s, drawn randomly from a uniform distribution.

Note that we varied the magnitude of reward (high vs. low), rather than presence vs. absence of reward. Hence, the task was designed to produce ceiling performance to ensure that participants would receive the high or low reward in the majority of trials. Trials in which participants did not gain the high or low reward were excluded from further analysis. Additionally, we presented the image and frame simultaneously to cue reward magnitude, rather than using a pre-stimulus reward cue. Reward probability was 100%, provided that participants correctly identified the reward magnitude during the choice phase of the trial. These last two measures were taken to ensure that participants paid attention to the image, and to shift the assumed dopamine response from the reward outcome to the reward cue presentation (image plus frame) (Shohamy and Adcock, 2010). Additionally, a fraction of the winnings (up to €5, with high reward items 20x more valuable than low reward items) was paid out directly after the fMRI session on day 1. This was done to ensure that participants would not discount reward magnitude due to delayed gratification (Peters and Büchel, 2011).

#### Day 2: Surprise Recognition Task (behavioural)

On the following day, participants returned to the lab to complete a surprise recognition memory test. All 120 objects and 120 scenes from the fMRI task (targets) were presented again without the coloured frames, together with 60 objects and 60 scenes that served as distractors. Stimuli were presented in 6 blocks with short resting breaks between blocks. Stimuli order was pseudo-randomised such that each block contained equal trial numbers of each condition, and no more than 3 stimuli belonging to the same level of each factor (category, reward), and no more than 3 distractors appeared in a row. For each image, participants indicated whether the image was ALT (“old”, presented during the fMRI task) or NEU (“new”). “Old” judgments were followed up by a choice between ERINNERT (“remember”), BEKANNT (“know”) or GERATEN (“guess”). This is an established procedure to distinguish between two processes thought to contribute to recognition memory: Recollection and familiarity (Tulving, 1985; Yonelinas et al., 2010). Participants were carefully instructed to only indicate “remember” if they had a vivid recollection of the image, including recall of contextual information. This was followed by two source memory tasks, consisting of forced-choice screens for the frame colour and for the reward magnitude. On the other hand, “new” judgments were followed up by a choice between SICHER (“sure”) and GERATEN (“guess”), without the frame and reward screens. All judgments were self-paced except the initial “old”/”new” judgment (4s).

#### Conditions of interest

In the encoding task, we manipulated item category (OB, object; SC, scene), and reward (HI, high; LO; low), resulting in 4 combinations: OB-HI, OB-LO, SC-HI, SC-LO. The “old”/”new” choices from the day 2 recognition phase were then used to back-sort the trials from the day 1 encoding phase into the following conditions of interest: OB-HI-H (object – high reward – hit), OB-HI-M (object – high reward – miss), OB-LO-H (object – low reward – hit), OB-LO-M (object – low reward – miss), SC-HI-H (scene – high reward – hit), SC-HI-M (scene – high reward – miss), SC-LO-H (scene – low reward – hit), and SC-LO-M (scene – low reward – miss).

### 2.3 Behavioural analyses

For the encoding task, we analysed the proportion of trials in which participants correctly identified the reward magnitude, calculated separately for each encoding condition (OB-HI, OB-LO, SC-HI, SC-LO). These proportions were then submitted to a two-way repeated measures ANOVA with the factors category and reward. For the recognition task, we calculated corrected recognition (CR) as the hit rate (proportion of “old” responses for targets) minus the false alarm rate (proportion of “old” responses to distractors, calculated separately for object and scene distractors). Additionally, from the distributions of “remember” and “know” responses, we calculated estimates for recollection and familiarity using the formula described in (Yonelinas and Jacoby, 1995). CR, recollection, and familiarity were calculated separately for each encoding condition (OB-HI, OB-LO, SC-HI, SC-LO) and submitted to two-way repeated measures ANOVAs with the factors category and reward. Similar analyses were conducted on the source memory responses for frame colour (source_frame_) and reward magnitude (source_reward_). As source memory involves retrieval of contextual detail, which is usually associated with recollection (Eichenbaum et al., 2007), we conducted source memory analyses only for recollected (“remember”) trials.

### 2.4 MRI acquisition

The study was scanned on a Siemens Tim Trio 3T MRI scanner using a 32-channel head coil. First, a high-resolution T1-weighted structural image was scanned (MPRAGE, 1mm isotropic voxels). Then, six functional runs were acquired using a T2*-weighted gradient-echo, echo-planar pulse sequence (40 interleaved slices, 1.5×1.5mm in-plane resolution, 2mm slice thickness with 20% distance factor, TR=1800ms, TE=30ms, multiband factor=2, PAT factor (GRAPPA)=2, 260 volumes per run). Slices were oriented in parallel to the AC-PC line and adjusted to optimise PFC coverage, with the field of view covering nearly the whole brain excepting very superior frontal and parietal cortex. The first 5 images of each functional run were discarded to allow for magnetic field stabilisation. Additionally, a 3D magnetisation transfer (MT) FLASH structural image was acquired (1mm isotropic voxels) after the functional runs.

### 2.5 fMRI preprocessing and analysis

#### Strategy

To account for the interindividual variability of MTL anatomy (Pruessner et al., 2002), our main analyses were carried out in individual space within bilateral regions of interest (ROIs). These encompassed the MTL subregions HC, PRC, and PHC, and additionally AMY and SNVTA (see Figure 2). The MTL and AMY ROIs were manually segmented on each participant’s T1 image using established landmarks (Insausti et al., 1998; Pruessner et al., 2000, 2002). Given previous findings that object and scene selectivity changes gradually along the MTL cortex axis (Litman et al., 2009; Liang et al., 2013), to optimise category selectivity we discarded the putative transition zone (posterior PRC and anterior PHC) in line with previous studies (Staresina et al., 2011, 2012, 2013; Schultz et al., 2019). The SNVTA ROI was manually segmented on each participant’s MT image as described in (Bunzeck and Düzel, 2006). In addition to the ROI analyses, control analyses were carried out on a voxel-wise level in Montreal Neurological Institute (MNI) space.

**Figure 2.**
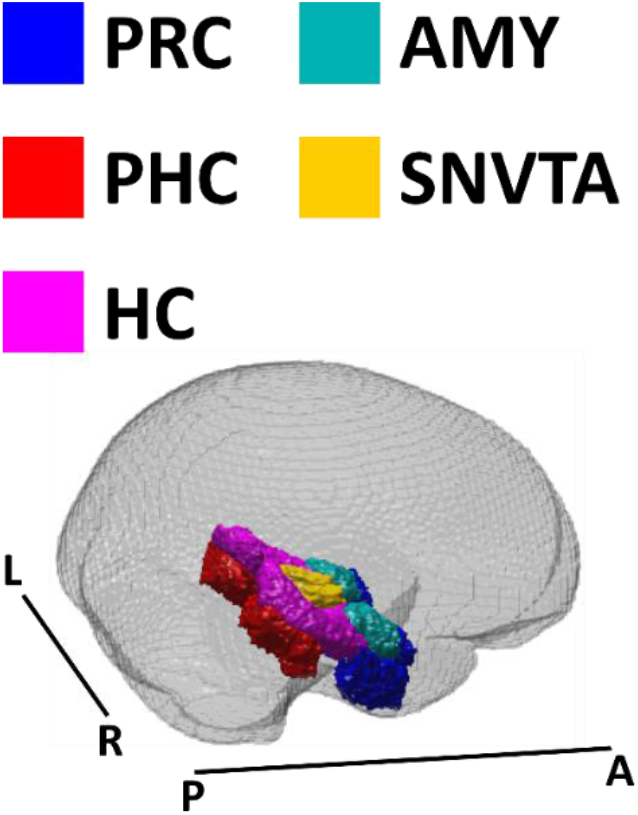
ROIs. Single-participant regions of interest were MNI-normalised, averaged over the full sample (n=39), and thresholded at 0.5. Here they are visualised within the standard SPM12 brain mask (mask_ICV.nii). Abbreviations: PRC: perirhinal cortex, PHC: parahippocampal cortex, HC: hippocampus, AMY: amygdala, SNVTA: substantia nigra/ventral tegmental area, A: anterior, P: posterior, R: right, L: left.

#### fMRI analysis pipeline

Functional runs were first corrected for differences in slice acquisition time, then realigned and unwarped to correct for movement and movement-related distortions using algorithms implemented in SPM12 (Wellcome Trust Centre for Neuroimaging, London, UK; http://www.fil.ion.ucl.ac.uk/spm/). The structural T1 image was coregistered to the mean functional volume using SPM12, and alignment was further improved using boundary-based registration as implemented in FSL epi_reg. The MT image was then coregistered to the T1 using SPM12. First-level statistical analyses (see below for details) were carried out on the non-normalised, unsmoothed data. For the ROI analyses, the ROIs were resampled to functional space, and first-level beta values were averaged across all voxels of each ROI. For the additional voxel-wise analyses, the T1 images were segmented into grey matter, white matter, and cerebrospinal fluid using SPM12. Deformation fields from this step were then used for MNI normalisation of the first-level beta images, and the normalised beta images were resampled to a 1mm isotropic voxel size, and smoothed with a Gaussian kernel (6mm full width at half maximum).

#### fMRI statistics

First-level general linear models were set up in SPM12. The six functional runs were concatenated. To account for this, the high-pass filter (128s) and autoregressive model AR(1) were adapted, and session constants included in the model. For our main analyses (model 1), the following conditions of interest were modelled: OB-HI-H, OB-HI-M, OB-LO-H, OB-LO-M, SC-HI-H, SC-HI-M, SC-LO-H, SC-LO-M. Conditions were modelled as impulse regressors using a canonical hemodynamic response function (HRF). Each trial phase (cue, choice, outcome) was modelled separately, and subsequent analyses were focused on the cue phase only. Additional regressors of no interest were included to model error trials, separately for object and scene trials. Error trials were defined as incorrect or no response during the Incidental Encoding Task, and/or no response during the old/new choice of the Recognition Task. Model 2 was set up identically, with the exception that the conditions of interest did not include the memory factor (hence, OB-HI, OB-LO, SC-HI, SC-LO). For the ROI analyses (model 1), the resulting beta images for each condition of interest were averaged across voxels of each participant’s ROIs, before being submitted to a group-level four-way repeated-measures ANOVA with the factors region, category, reward, and subsequent memory. Where appropriate, Greenhouse-Geisser correction was applied. Follow-up analyses were then carried out within each ROI. For the voxel-wise analyses (model 2), normalised, smoothed beta maps were submitted to a second-level random effects analysis (flexible factorial as implemented in SPM12) that included the factors category and reward as well as a subject factor. The resulting brain activation maps were corrected for multiple comparisons using peak-level family-wise error correction within a study-specific MNI brain mask consisting of the following: (i) the manually delineated masks of HC and SNVTA, normalized to MNI space and averaged over the full sample (n=39), thresholded at 0.5, and (ii) an existing mask from the Rangel Neuroeconomics Laboratory (www.rnl.caltech.edu/resources/index.html), which contains brain regions consistently implicated in reward processing including vmPFC, VS, and posterior cingulate cortex (PCC) (Bartra et al., 2013; Clithero and Rangel, 2014).

## 3 Results

### 3.1 Behavioural results

#### Incidental encoding task

As expected, accuracy in the incidental encoding task was near ceiling (mean [SEM] % accuracy: OB-HI: 98.5 [0.4], OB-LO: 98.4 [0.4], SC-HI: 97.0 [0.7], SC-LO: 97.8 [0.5]). A repeated measures ANOVA with the factors category and reward showed a significant effect of category (higher accuracy for object trials, *F*_(1,26)_=5.111, *p*=.032), but no effect of reward or interaction of category and reward (*p*≥.212). Importantly, only trials with accurate responses in the encoding task were considered in the behavioural analyses of recognition memory as well as the fMRI analyses.

#### Recognition task

For the recognition task, we expected improved subsequent memory for high-reward compared to low-reward items for both objects and scenes (see Table 1 for overview). We analysed corrected recognition (CR, hit rate minus false alarm rate) for each condition. A repeated measures ANOVA yielded a main effect of reward (high > low, *F*_(1,26)_=18.297, *p*<.001) and, unexpectedly, a main effect of category (objects > scenes, *F*_(1,26)_=7.404, *p*=.011) as well as an interaction effect of category and reward (*F*_(1,26)_=9.961, *p*=.004). Follow-up paired t-tests indicated that high-reward objects were remembered better than low-reward objects (OB-HI>OB-LO, *t*_(26)_=5.568, *p* <.001), while high-reward scenes were remembered better than low-reward scenes on a trend level only (SC-HI>SC-LO, *t*_(26)_=1.759, *p*=.090). Hence, the observed interaction effect indicates a greater reward enhancement of object memory compared to scene memory. We also explored whether these results reflected a general difference between objects and scenes, e.g. due to systematic differences in visibility or memorability between the two categories. Such a difference should be apparent in both the high-reward and low-reward condition. However, the difference between objects and scenes was only significant in the high-reward (OB-HI vs. SC-HI, *t*_(26)_=3.590, *p*=.001), but not in the low-reward condition (OB-LO vs. SC-LO, *t*_(26)_=0.596, *p*=.556), indicating that the observed main effect of category was driven by the interaction effect.

**Table 1.** Overview over recognition memory results, demonstrating consistent effects of our experimental manipulations across all outcome measures at both sample sizes. The table contains mean (SEM) values for all four conditions as well as *F* and *p* values from two-way repeated measures ANOVAs with the factors category and reward. ^1^Memory sample (n=27), ^2^full sample (n=39).

| Outcome | OB-HI | OB-LO | SC-HI | SC-LO | Effect of category | Effect of reward | Interaction |
| --- | --- | --- | --- | --- | --- | --- | --- |
| CR<sup>1</sup> | 0.382<br>(0.023) | 0.295<br>(0.017) | 0.317<br>(0.020) | 0.287<br>(0.018) | $F_{(1,26)}=7.404$,<br>$p=0.011$ | $F_{(1,26)}=18.297$,<br>$p<0.001$ | $F_{(1,26)}=9.961$,<br>$p=0.004$ |
| CR<sup>2</sup> | 0.330<br>(0.022) | 0.222<br>(0.023) | 0.254<br>(0.023) | 0.231<br>(0.020) | $F_{(1,38)}=4.386$,<br>$p=0.043$ | $F_{(1,38)}=29.980$,<br>$p<0.001$ | $F_{(1,38)}=26.278$,<br>$p<0.001$ |
| Recollection<sup>1</sup> | 0.194<br>(0.021) | 0.124<br>(0.015) | 0.138<br>(0.021) | 0.117<br>(0.018) | $F_{(1,26)}=3.209$,<br>$p=0.085$ | $F_{(1,26)}=18.711$,<br>$p<0.001$ | $F_{(1,26)}=10.018$,<br>$p=0.004$ |
| Recollection<sup>2</sup> | 0.173<br>(0.017) | 0.103<br>(0.012) | 0.118<br>(0.016) | 0.106<br>(0.014) | $F_{(1,38)}=3.903$,<br>$p=0.055$ | $F_{(1,38)}=19.582$,<br>$p<0.001$ | $F_{(1,38)}=13.536$,<br>$p<0.001$ |
| Familiarity<sup>1</sup> | 1.016<br>(0.092) | 0.800<br>(0.075) | 0.836<br>(0.077) | 0.777<br>(0.072) | $F_{(1,26)}=1.949$,<br>$p=0.175$ | $F_{(1,26)}=9.482$,<br>$p=0.005$ | $F_{(1,26)}=6.372$,<br>$p=0.018$ |
| Familiarity<sup>2</sup> | 0.868<br>(0.078) | 0.598<br>(0.077) | 0.663<br>(0.074) | 0.584<br>(0.071) | $F_{(1,38)}=3.376$,<br>$p=0.074$ | $F_{(1,38)}=19.642$,<br>$p<0.001$ | $F_{(1,38)}=14.603$,<br>$p<0.001$ |

Additional analyses were conducted to explore whether the observed memory effects were specific to a memory process (recollection or familiarity, see Materials and Methods) or sample (memory subsample as in the analyses above, n=27, or full sample, n=39). Importantly, all memory measures (CR, recollection, familiarity) at both sample sizes showed the observed interaction between category and reward in the same direction, with greater reward enhancement of object memory than scene memory (memory subsample: *F*_(1,26)_≥9.961, *p*≤.004; full sample: *F*_(1,38)_ ≥6.372, *p*≤.018, see Table 1).

#### Source memory

Finally, we analysed the responses to the source memory tasks, in which participants indicated which frame colour (source_frame_) and reward magnitude (source_reward_) had been associated with each image. Note that source memory analyses were conducted for “remember” trials only, which reduced the memory sample to 19 participants with at least four “remember” responses per condition, or fewer than half of the original sample size. For source_frame_, we analysed the proportion of correct responses (mean [SEM] source_frame_: OB-HI: 0.587 [0.031], OB-LO: 0.474 [0.026], SC-HI: 0.578 [0.044], SC-LO: 0.494 [0.044]). Source_frame_ exceeded chance performance for OB-HI (*t*_(18)_=2.774, *p*=.013) and, on a trend level, for SC-HI (*t*_(18)_=1.766, *p*=.094), but not for OB-LO or SC-LO (all *p*≥0.340). A repeated measures ANOVA with the factors category and reward yielded a main effect of reward (high > low, *F*_(1,18)_=11.497, *p*=.003, all other *p*≥0.675). Paired t-tests revealed a significant difference between OB-HI and OB-LO (*t*_(18)_=3.307, *p*=.004), but not between SC-HI and SC-LO (*p*=.137). For source_reward_, we analysed the proportion of trials in which participants indicated that an image had been paired with high reward (mean [SEM] source_reward_: OB-HI: 0.765 [0.041], OB-LO: 0.689 [0.052], SC-HI: 0.737 [0.040], SC-LO: 0.633 [0.056]). Note that, as high memory confidence may in itself be rewarding (Schwarze et al., 2013), these trials are biased towards “high reward” source responses in all conditions, and chance level is therefore meaningless. A repeated measures ANOVA with the factors category and reward yielded a main effect of reward (high > low, *F*_(1,18)_=10.962, *p*=.004, all other *p*≥0.234) such that participants were more likely to indicate “high reward” to high-reward than to low-reward items. Paired t-tests indicate that this was the case for both OB-HI vs. OB-LO (*t*_(18)_=2.432, *p*=.026) and SC-HI vs. SC-LO (*t*_(18)_=2.110, *p*=.049).

### 3.2 fMRI: ROI results – Category, subsequent memory, and the role of reward

#### Overall analysis

First, we analysed whether reward modulated memory encoding for objects and scenes in our ROIs (model 1, memory subsample). Beta values from our conditions of interest (OB-HI-H, OB-HI-M, OB-LO-H, OB-LO-M, SC-HI-H, SC-HI-M, SC-LO-H, SC-LO-M; with H: hit, M: miss in the subsequent recognition phase) were averaged across all voxels of each ROI (HC, PRC, PHC, AMY, SNVTA) and submitted to a four-way repeated-measures ANOVA with the factors ROI, category, reward, and subsequent memory. We report interactions of the ROI factor with any experimental factors. This analysis revealed significant two-way interactions of ROI with category (*F*_(1. 75,45.50)_= 133.036, *p*<.001) and memory (*F*_(2.85,74.02)_=9.819, *p*<.001), a three-way interaction of ROI with category and memory (*F*_(2.26,58.75)_=6.493, *p*<.001), and a four-way interaction of ROI with category, reward, and memory (*F*_(2.26,58.75)_=3.228, *p*=.020). The interaction of ROI with reward was marginally significant (*F*_(2.26,58.75)_=2.880, *p*=.051). There was no other interaction effects involving the ROI factor (all *p*≥.511).

Given the significant four-way interaction of ROI, category, reward, and memory, we computed individual three-way ANOVAs within each ROI as well as follow-up tests where appropriate. A summary of results for each ROI is given in Figure 3A.

**Figure 3.**
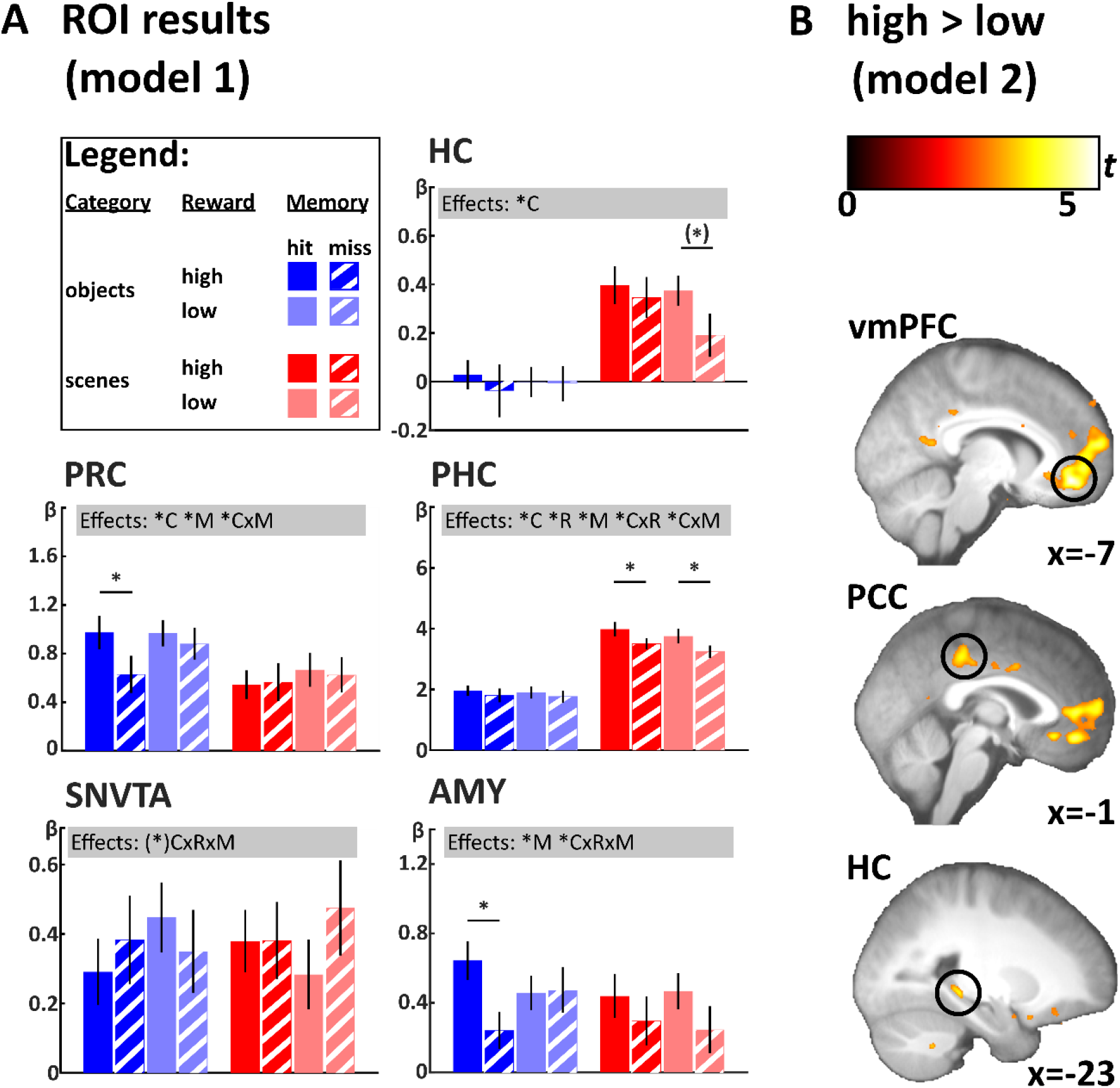
Overview over fMRI results. A. Average beta values for each condition of model 1 (memory sample, n=27) in each of the 5 ROIs. Grey boxes indicate results from individual three-way repeated measures ANOVA with the factors category (C), reward (R), and subsequent memory (M). \**p*<.05, (*)*p*<.1. Error bars indicate SEM. B. Voxel-wise activity for the high > low reward contrast in model 2 (full sample, n=39). Display threshold *p*<.001, k≥5 voxels. Statistical maps are projected onto the normalised, averaged T1.

#### HC

The HC showed a significant main effect of category (scenes > objects, *F*_(1,26)_=70.192, *p*<.001). No other main effect or interaction was significant (all *p*≥.207).

#### PRC

The PRC showed significant main effects of category (objects > scenes, *F*_(1,26)_=10.557, *p*=.003) and subsequent memory (hits > misses, *F*_(1,26)_=5.840, *p*=.023). Importantly, the interaction of category and subsequent memory was also significant (*F*_(1,26)_=6.558, *p*=.017), indicating that subsequent memory effects were stronger for objects than scenes. No other main or interaction effect was significant (all *p*≥.118). To explore which conditions showed subsequent memory effects, we additionally computed paired t-tests between subsequent hits and misses for OB-HI, OB-LO, SC-HI, and SC-LO. Notably, only the subsequent memory effect for OB-HI was significant (*t*_(26)_=3.036, *p*=.005, all other *p*≥.412).

#### PHC

The PHC showed significant main effects of category (scenes > objects, *F*_(1,26)_=153.697, *p*<.001) and subsequent memory (hits > misses, *F*_(1,26)_=29.407, *p*<.001). Importantly, the interaction of category and subsequent memory was also significant (*F*_(1,26)_=6.372, *p*=.018), indicating that subsequent memory effects were stronger for scenes than objects. Additionally, we observed a significant main effect of reward (high > low, *F*_(1,26)_=10.391, *p*=.003), as well as a significant interaction effect of category and reward (greater reward effect for scenes than objects, *F*_(1,26)_=4.658, *p*=.040) – note however the lack of a significant interaction of these effects with the ROI factor in the overall analysis above. Again, we explored which conditions showed subsequent memory effects using pairwise t-tests between subsequent hits and misses. We observed significant effects of subsequent memory for both SC-HI and SC-LO (*t*_(26)_=[3.788 3.998], *p*<.001), but not for either OB-HI or OB-LO (all *p* ≥.181).

#### AMY

The AMY showed a significant main effect of subsequent memory (hits > misses, *F*_(1,26)_=7.040, *p*=.013) as well as a significant three-way interaction of category, reward, and subsequent memory (*F*_(1,26)_=4.369, *p*=.047). To identify the constituents of this three-way interaction, we computed separate two-way ANOVAs (reward, memory) for objects and scenes, respectively, assessing the interaction effects only. AMY showed a significant interaction effect of reward and subsequent memory for objects (*F*_(1,26)_=7.183, *p*=.013) but not scenes (*p*=.651). Again, we explored which conditions showed subsequent memory effects using pairwise t-tests between subsequent hits and misses. Notably, only the subsequent memory effect for OB-HI was significant (*t*_(26)_=3.385, *p*=.002, all other *p*≥.101).

#### SNVTA

Contrary to our expectations, the SNVTA showed no significant main effects or interactions, save for a trend-level three-way interaction of category, reward, and subsequent memory (*F*_(1,26)_=2.957, *p*=.097, all other *p*≥.315). In particular, neither the main effect of reward nor the interaction of reward and subsequent memory were significant (all *p*≥.651).

### 3.3 fMRI: Voxel-wise effects of reward (model 2)

The previous sections demonstrated clear effects of our reward manipulation on behavioural measures of memory, for objects more so than scenes. The neural effects of reward on memory formation showed a similar asymmetry. Specifically, subsequent memory effects for objects in PRC and AMY were only significant for high-reward objects, while subsequent memory effects for scenes in PHC were significant for both high-reward and low-reward scenes. Against our expectations, however, we did not observe main effects of reward, or interaction effects of reward with subsequent memory that were independent of category, in either HC or SNVTA. Therefore, as a control analysis, we tested whether our task succeeded in engaging the reward network, and whether these effects differed between objects and scenes. We used a reduced model with two factors (category, reward) and tested for main effects of reward as well as interaction effects of reward and category. By disregarding the memory factor, we made use of the increased experimental power of the full sample (n=39). Additionally, we used a voxel-wise approach in MNI-normalised data to be able to identify small clusters of activity, which may not be picked up in an ROI analysis. Effects are reported for, and corrected for multiple comparisons within, a brain mask comprising vmPFC, VS, PCC, HC, and SNVTA (see Materials and Methods for details).

Table 2 gives an overview of the observed voxel-wise effects located inside the mask. The high > low reward contrast revealed clusters of activity with peaks in vmPFC (MNI [x y z]: [−7 42 −12], *t*_(38)_ = 5.74, *p*_FWE_<.001), PCC ([−1 −32 42], *t*_(38)_=4.4, *p*_FWE_=.024), and left posterior HC ([−23 −34 −5], *t*_(38)_=4.40, *p*_FWE_=.037). Additional clusters in bilateral HC, VS, and vmPFC emerged at an uncorrected threshold of *p*<.001. Notably, there was no activity in the SNVTA, even at a relaxed uncorrected threshold of *p*<.01. The objects x reward interaction ([OB-HI>OB-LO]>[SC-HI>SC-LO]) yielded clusters in vmPFC only at an uncorrected threshold of *p*<.001 (all *p*_FWE_≥.199). For the scene x reward interaction ([SC-HI>SC-LO]>[OB-HI>OB-LO]), a cluster in right HC was marginally significant ([30 −32 −2], *t*_(38)_=4.11, *p*_FWE_=.061). Additionally, clusters in right HC and SNVTA emerged at an uncorrected threshold of *p*<.001 (see Table 2).

**Table 2.** Overview of significant clusters from the voxel-wise analysis within a brain mask comprising vmPFC, VS, PCC, HC, and SNVTA (see Materials and Methods). Uncorrected threshold *p*<.001, cluster threshold: 5 voxels total including voxels outside mask. # voxels refers to the number of voxels within the mask. *p*_FWE_ refers to the *p* value of the peak voxel after family-wise error correction within the brain mask. ^1^For the inclusive masking analysis, we report coordinates and statistical values for OB-HI>OB-LO inclusively masked with SC-HI>SC-LO (outside brackets), and vice versa (inside brackets). \**p*_FWE_<.05, (*)*p*_FWE_<.1.

| Contrast | Region | # voxels | x | y | z | t | $p_{FWE}$ |
| --- | --- | --- | --- | --- | --- | --- | --- |
| Main effect: High > low | vmPFC | **5063** | **-7** | **42** | **-12** | **5.74** | **<.001\*** |
|  |  | 8 | -25 | 28 | -17 | 3.65 | .232 |
|  | PCC | **416** | **-1** | **-32** | **42** | **4.40** | **.024\*** |
|  | L HC | **189** | **-23** | **-34** | **-5** | **4.27** | **.037\*** |
|  |  | 79 | -33 | -27 | -16 | 3.75 | .178 |
|  |  | 16 | -32 | -13 | -19 | 3.61 | .257 |
|  |  | 16 | -17 | -15 | -26 | 3.59 | .271 |
|  | R HC | 109 | 20 | -29 | -13 | 3.90 | .116 |
|  |  | 66 | 26 | -18 | -21 | 3.84 | .141 |
|  |  | 23 | 32 | -16 | -15 | 3.43 | .383 |
|  | VS | 1 | 5 | 10 | -1 | 3.70 | .204 |
|  |  |  | 2 | 12 | -1 | 3.27 | .525 |
| Interaction:<br>(OB-HI > OB-LO) ><br>(SC-HI > SC-LO) | vmPFC | 47 | -5 | 25 | -14 | 3.71 | .199 |
|  |  | 5 | -6 | 26 | -11 | 3.54 | .304 |
|  |  | 28 | 0 | 38 | -12 | 3.45 | .369 |
|  |  | 1 | -2 | 32 | -23 | 3.19 | .598 |
| Interaction:<br>(SC-HI > SC-LO) ><br>(OB-HI > OB-LO) | R HC | **54** | **30** | **-32** | **-2** | **4.11** | **.061(\*)** |
|  |  | 28 | 35 | -22 | -14 | 3.54 | .304 |
|  | SNVTA | 16 | 3 | -24 | -20 | 3.32 | .476 |
| Inclusive masking<sup>1</sup>:<br>OB-HI>OB-LO and<br>SC-HI>SC-LO | vmPFC | 8 | -7 (-7) | 46 (47) | -15 (-15) | 4.27 (3.29) | .038 (.509) |
|  |  | 48 | -8 (-8) | 50 (50) | -5 (-3) | 3.97 (3.42) | .095 (.395) |
|  |  | 24 | -7 (-7) | 41 (41) | -6 (-4) | 3.75 (3.39) | .181 (.419) |
|  |  | 26 | -1 (0) | 44 (43) | 4 (4) | 3.73 (3.60) | .187 (.264) |
|  |  | 1 | -2 (-2) | 52 (52) | 6 (6) | 3.47 (3.17) | .353 (.620) |
|  |  | 4 | -4 (-6) | 54 (54) | 4 (4) | 3.44 (3.22) | .376 (.570) |

Lastly, as main effects in a factorial design may be driven by interaction effects, we identified brain regions in which both the OB-HI > OB-LO and SC-HI > SC-LO contrast exceeded a threshold of *p*<.001 uncorrected (inclusive masking approach). This analysis yielded clusters located in the vmPFC (see Table 2).

## 4 Discussion

### Summary

The present study’s goals were twofold: One, to investigate whether the documented dichotomy between PRC and PHC for object and scene memory extends to item encoding, and two, to investigate whether reward modulates item encoding of objects and scenes. Behaviourally, high reward predominantly enhanced subsequent memory for objects in both model-free (CR) and modelbased (recollection, familiarity) outcome measures. Evidence for reward enhancement of memory for scenes, on the other hand, was modest. Importantly, neural activity in PRC and PHC predicted subsequent item memory for objects and scenes, respectively. Furthermore, neural encoding activity exhibited an asymmetry that mirrored our behavioural findings: Encoding activity in AMY was selective for high-reward objects, with a similar (albeit non-significant) pattern in PRC, while encoding activity in PHC did not differ between high- and low-reward scenes. Finally, reward-related brain activity, regardless of stimulus category, was centred on the vmPFC.

### Category-specific incidental item encoding in the MTL cortex

We observed opposite patterns of category-specific incidental item encoding in PRC and PHC, with object encoding in PRC and scene encoding in PHC. Category specificity as an organising principle for the functional architecture of the MTL has been shown in a number of imaging studies for processes including perception (Litman et al., 2009; Liang et al., 2013; Berron et al., 2018), associative encoding (Awipi and Davachi, 2008; Staresina et al., 2011), associative retrieval (Staresina et al., 2012, 2013; Mack and Preston, 2016; Schultz et al., 2019), working memory (Libby et al., 2014), short-term memory reactivation (Schultz et al., 2012), and recognition memory (Martin et al., 2013; Kafkas et al., 2017). However, to our knowledge, our study is the first to demonstrate a double dissociation between PRC and PHC for category-specific item encoding, thereby filling an important gap in the literature and supporting a model of MTL function that draws on anatomical connectivity to predict functional specialisation (Davachi, 2006; Eichenbaum et al., 2007).

While scene-specific item encoding has been previously observed in PHC and the larger parahippocampal place area (Prince et al., 2009; Preston et al., 2010), the same cannot be said for object-specific item encoding in PRC. In their 2010 paper, Preston and colleagues investigated incidental encoding of faces and scenes during a target detection task, observing subsequent memory effects for scenes in PHC, and subsequent memory effects for both faces and scenes in PRC (note that restricting the analysis to face-selective voxels in PRC yielded a response pattern consistent with face-specific encoding, however, the interaction between category and memory was not significant). In contrast, PRC in our study did not show a subsequent memory effect for scenes at all. Preston and colleagues’ study design differs from ours in a number of ways. They assessed recognition memory immediately after the encoding task rather than after a one-day delay, and their participants were aware that they would be tested on their recognition memory as the study included two encoding-recognition cycles. Potentially, intentional encoding leads to increased elaboration of the encoding stimuli and therefore to additional recruitment of PRC during scene encoding (however, memory performance [CR, hits minus false alarms] is roughly matched between the two studies). Additionally, we note that on a descriptive level, our object encoding effect in PRC appears to be specific to high-reward objects. It is possible that while incidental encoding does not engage PRC in a category-specific fashion, adding a motivational factor such as reward does. We will return to this line of thought later in the discussion.

We observed category specificity in the MTL cortex during item encoding. Previous work has demonstrated such effects for associative encoding (Awipi and Davachi, 2008; Staresina et al., 2011). It has been argued (Davachi, 2006; Eichenbaum et al., 2007) that item memory is a distinct process from associative, relational, or context memory. In this view, item memory stems from object-related processing in the anterior MTL cortex (PRC), and is associated with the subjective sensation of familiarity, whereas associative memory is related to spatial and multi-modal processing in the posterior MTL cortex (PHC) and HC, and associated with the subjective sensation of recollection. Nevertheless, we observed scene-specific item memory in the PHC. Notably, the present results as well as prior work (Awipi and Davachi, 2008; Staresina et al., 2011, 2012, 2013; Martin et al., 2013; Schultz et al., 2019) suggest that category specificity in the MTL cortex emerges even if memory processes are held constant. Thus, category specificity is an essential commonality between different memory processes, even if object and spatial processing may map preferentially onto item and associative/ recollective memory, respectively (Davachi, 2006; Eichenbaum et al., 2007). However, PRC and PHC also distinguish between item and context recall even when the stimulus material is being held constant (Wang et al., 2013), and studies have shown dissociable effects of experimental manipulations on different memory processes (Wittmann et al., 2005, 2011; Bisby and Burgess, 2013; McCullough et al., 2015; Madan et al., 2017; Ritchey et al., 2019). This implies that process dissociations may have additional predictive value for MTL function that go beyond a distinction based on stimulus categories. Future research may determine how exactly category specificity maps onto process dissociations.

### Asymmetric effects of reward on behavioural and neural measures of memory

We observed a preferential enhancement of object memory by reward, accompanied by a similar asymmetry in the neural data: Subsequent memory effects in AMY were selective for high-reward objects, with a similar, albeit non-significant, pattern in PRC, while PHC showed subsequent memory effects for both high- and low-reward scenes. (Note that this effect is unlikely to reflect a general difference between object and scene encoding, for example due to differences in salience or luminance between the categories, since memory for low-reward objects did not differ from memory for low-reward scenes.) The clear behavioural bias for high-reward objects in the present study is surprising, given that previous studies have shown motivational effects on encoding when only scenes were used (Adcock et al., 2006; Bunzeck et al., 2012; Spaniol et al., 2014; Rouhani et al., 2018) but see e.g. (Steiger and Bunzeck, 2017). One potential reason for this could be memory performance. The incidental encoding task in the present study yielded a comparably low memory performance that was based mainly on familiarity rather than recollection, while reward effects on memory have been associated with recollection or high-confidence hits (Wittmann et al., 2005, 2011; Adcock et al., 2006). It is therefore possible that a more robust reward effect on scenes would have emerged with higher proportions of recollection.

More intriguingly, the observed behavioural and neural asymmetry may be explained within an existing framework of memory, the PMAT framework (Ranganath and Ritchey, 2012; Ritchey et al., 2015). This theory poses that two large-scale brain networks underlie memory, with an anterior-temporal (AT) system, including PRC and AMY, representing objects and their motivational significance, and a posterior-medial (PM) system, including PHC, representing (spatial) context. Indeed, PRC and AMY have been associated with acquiring stimulus-reward associations (Liu and Richmond, 2000; Liu et al., 2000; Rudebeck et al., 2017). In this view, the HC may play a role both in sharpening and integrating information received from these systems (Ranganath and Ritchey, 2012; Ritchey et al., 2015). Previously, the HC, in interplay with SNVTA, has also been implicated in reward enhancement of episodic encoding of highly confident or recollected items (Wittmann et al., 2005; Adcock et al., 2006). Here, in a task that yielded low memory performance based mainly on familiarity, we observed a behavioural advantage of high-reward object encoding and subsequent-memory effects for high-reward objects in two putative AT regions, AMY and PRC, but not in HC or SNVTA. It is possible that the AT system is sufficient for supporting reward enhancement of low-confident object memory. But if HC integrates information from both streams, then perhaps item memory for scenes, processed preferably along regions of the PM system, is less likely to receive an advantage from motivational factors unless it reaches HC’s recollection threshold. Indeed, we are not aware of studies showing motivational enhancement of low-confident scene memory, and it is worth pointing out that our source memory effects, computed for recollected trials only, appear more balanced between objects and scenes. These considerations are somewhat speculative and require further research. One prediction would be that the observed asymmetry between reward enhancement of objects and scenes decreases at higher rates of highly confident memory responses.

In a recent study, Ritchey and colleagues (2019) found that PRC and AMY supported encoding of emotional over neutral items, whereas the PHC and HC supported context encoding for both emotional and neutral items, but were not engaged in item encoding. Behavioural data showed a similar asymmetry: Emotion enhanced item memory but did not affect context memory. Item and context processing in the MTL cortices have been previously linked to their putative roles in object-related and spatial processing (Davachi 2006, Eichenbaum 2007), facilitating parallels between Ritchey et al.’s results and ours. Modulation of item encoding by motivational factors such as emotion or, in our case, reward, may not require the HC, but instead be carried by PRC and AMY (note, however, that in this study, the items consisted of scene images, and the context consisted of tasks solved during encoding).

### Category specificity and category independence in the HC

The HC showed a robust effect of scene viewing compared to object viewing. While some accounts of MTL see the HC’s role in memory as category-independent (Davachi, 2006; Eichenbaum et al., 2007), others emphasise the role of HC in spatial (Moser et al., 2008; Hartley et al., 2014) and scene processing (Maguire and Mullally, 2013). Moreover, while we observed a pronounced scene effect in HC in the present dataset, an earlier study – using a similar stimulus set, albeit a different (intentional, associative) encoding task – did not (Schultz et al., 2019). Recent work indicates that different subfields of HC may be differentially sensitive to both task and stimuli (Dalton et al., 2018). Targeted high-resolution investigations of HC subfields may further specify the circumstances in which HC responses are category-specific or category-independent. We also did not observe overall effects of subsequent memory in HC. HC has been implicated in recollective, or highly confident recognition memory (Wittmann et al., 2005; Adcock et al., 2006; Eichenbaum et al., 2007). Memory performance in the present study was overall low, which may explain why neural encoding processes were mainly observed in the MTL cortex, thought to support familiarity (Eichenbaum et al., 2007; Martin et al., 2013), rather than HC.

### Reward-related processing in the MTL and beyond

Against our hypotheses, we did not observe reward responses in the SNVTA and only limited evidence for reward processing in the HC and VS. Activity in the SNVTA and VS, a major target region of the SNVTA’s dopaminergic projections (Haber and Knutson, 2010), has been shown to vary with prediction error (Schultz, 1998; O’Doherty et al., 2003; Pessiglione et al., 2006; D’Ardenne et al., 2008; Rolls et al., 2008). Hence, the task developed for the present study aimed to shift the prediction error and therefore the putative dopaminergic response to the presentation of the encoding cue (see Materials and Methods). It is, however, not a learning task, as the reward contingencies were explicitly conveyed to the participants prior to each run. Diederen and colleagues (Diederen et al., 2016) argued that SNVTA’s role in adaptive prediction error coding may be particularly pronounced in tasks that require learning of reward contingencies. While previous studies have shown SNVTA engagement during encoding of stimuli in tasks that did not require such learning (Wittmann et al., 2005; Adcock et al., 2006), future work may show whether SNVTA/VS prediction error signalling in a reinforcement learning task covaries with successful episodic memory encoding. So far, it has been demonstrated that encoding of *irrelevant* objects during cue presentation actually interferes with striatal learning (Wimmer et al., 2014), but it is unclear whether the same holds true for episodic encoding of the reward-predicting cues.

On the other hand, we observed robust reward signals in the vmPFC. The vmPFC is a major part of the brain’s reward system (Haber and Knutson, 2010), and thought to code a range of processes, including subjective value (Kable and Glimcher, 2009; Hebscher and Gilboa, 2016). Intriguingly, it is also thought to play a role in memory, namely the acquisition and utilisation of abstract knowledge structures, so-called schemas (Hebscher and Gilboa, 2016). In our encoding task, participants had to match a stimulus (e.g. an image of a coffee cup surrounded by a blue frame) to an existing abstract knowledge structure (e.g. “objects surrounded by a blue frame signal high reward”). Memory schemas predicting high reward may be encoded more deeply than schemas predicting low reward, leading to elevated vmPFC engagement. Indeed, recent work implies that vmPFC is necessary for processing configural objects in which a combination of features, but not one feature alone, signals their value (Pelletier and Fellows, 2019). Similarly, in the present task, neither stimulus category nor frame colour, but only their combination, signalled reward magnitude. Effects of high reward were observed in vmPFC for both object and scene trials. While the vmPFC has been suggested as a convergence zone of the AT and PM systems (Ranganath and Ritchey, 2012; Ritchey et al., 2015), anatomical connectivity between that region and the MTL cortex varies along the anterior-posterior MTL axis (Kondo et al., 2005; Price, 2007; Kondo and Witter, 2014) and may be particularly pronounced for anterior MTL cortex (PRC, entorhinal cortex) (Eichenbaum, 2017). However, anatomical labelling may not be directly comparable between species (Price, 2007; Haber and Knutson, 2010), and resting-state connectivity in humans has indicated preferential functional connectivity of the vmPFC with PHC rather than PRC (Kahn et al., 2008). Therefore, a query for future work is whether vmPFC engagement during a reward task could bias reward-related object encoding by modulating one MTL pathway over the other.

### Future directions

The fate of a memory trace is not solely determined by neural processing during encoding. For example, reward during encoding may enhance post-encoding consolidation processes, by biasing recently-encoded memories for offline replay (Kumaran et al., 2016). Similarly, reward associations acquired during encoding modulate brain activity during retrieval (Wolosin et al., 2012; Elward et al., 2015). In sum, the observed behavioural effects of reward on object memory may have been driven in part by neural processing outside the time window observed in the present study, which may be addressed in future work.

### Conclusions

In sum, we present novel evidence for a double dissociation between anterior and posterior MTL cortex for incidental item encoding of objects and scenes, respectively. Additionally, reward preferably modulated object rather than scene encoding, evident in behavioural measures and anterior temporal lobe signalling. A potential limitation of our study lies in the comparatively low memory performance, which was mainly based on familiarity rather than recollection. Future work may further elucidate the neural mechanisms underlying the distinct effects of reward on the encoding of different stimulus categories.

## Notes

**Conflict of interest:** The authors declare no conflict of interest.

### Competing Interest Statement

The authors have declared no competing interest.

